# Exercise-related genes analysis of Mongolian Horse

**DOI:** 10.1101/327536

**Authors:** Jing Pan, Chimge Purev, Hongwei Zhao, Zhipeng Zhang, Feng Wang, Nashun Wendoule, Guichun Qi, Huanmin Zhou

**Affiliations:** College of Life Sciences, Inner Mongolia Agricultural University, Huhehot, China; Mongolia-China Joint Laboratory of Applied Molecular Biology, “Administration of the Science Park” CSTI, Ulaanbaatar, Mongolia; Beijing 8omics Gene Technology Co.Ltd, Beijing, China; College of Life Sciences, Nankai University, Tianjin, China; Animal Husbandr Workstation of Ewenki Autonomous County, Hulun Buir, China; Bayanta Village of Animal Husbandry and Veterinary Station of Ewenki Autonomous County, Hulun Buir, China

**Keywords:** genome, Mongolian horse, Abaga horse, Wushen horse, exercise

## Abstract

The Mongolian horses, as a neglected scientific resource, have excellent endurance and stress resistance to adapt to the cold and harsh plateau conditions. Intraspecific genetic diversity is mainly embodied in various genetic advantages of different branches of Mongolian horse. Abaga horse is better than Wushen horse in running speed, for example. Because people pay progressively attention to the athletic performance of horse, such as horse racing in Mongolia’s Naadam festival, we expect to guide the exercise-oriented breeding of horses through genomics research. We obtained the clean data of 630,535,376,400 bp through the entire genome second-generation sequencing for the whole blood of 4 Abaga horses and 10 Wushen horses. Based on the data analysis of single nucleotide polymorphism (SNP), we severally detected that 479 and 943 positively selected genes, particularly exercise-related, were mainly enriched on equine chromosome 4 in Abaga horses and Wushen horses, which implied that the chromosome 4 may be associated with the evolution of the Mongolian horse and athletic performance. Four hundred and forty genes of positive selection were enriched in 12 exercise-related pathways and narrowed in 21 exercise-related genes in Abaga horse, which were distinguished from Wushen horse. So, we speculated that the Abaga horse may have oriented genes for the motorial mechanism and 21 exercise-related genes also provided molecular genetic basis for exercise-directed breeding of Mongolian horse.

## INTRODUCTION

As an ancient breed, Mongolian horse has gone through a long breeding period (China National Commission of Animal Genetic Resources 2011). With a view to research tendentiousness, researchers pay more attention to traits of Thoroughbred horse (Gim *et al.* 2004; Park *et al.* 2012; Capomaccio *et al.* 2013) and Quarter horse (Doan *et al.* 2012; Meira *et al.* 2014), but not the Mongolian horse and its diverse sub-branch. The preeminent endurance and stress-resistance of Mongolia horse are important factors for it to well adapt to the cold and harsh plateau environment (Li *et al.* 2009). Natural factors may have enormous impacts on evolution owing to the rough domestication of Mongolian horse (Hund 2008). Due to various geographic conditions and human necessities, Mongolian horse gradually formed several specific traits. Some horses which adapt to the desert climate have larger feet, for instance, some horses which adjust to mountain road with rocks have supple body and hard hoof; in addition, the features of horse which accommodate to the grassland climate are tall physique and good at running (Elisabeth 2011). Living in the Xilin Gol grassland of Inner Mongolia, Abaga horse belongs to the steppe horse and speeds up to 1600 meters every 91.47 seconds (China National Commission of Animal Genetic Resources 2011). Wushen horse, which is small build and has broad-flat horseshoe, as symbol of the desert horse in the south of Maowusu desert of Ordos City in Inner Mongolia, can hoof steadily in the desert, albeit noffast running speed of 13 to 15 kilometers per hour (Dugarjaviin 2009).

In Mongolia, the herdsmen depend on horses by reason that they are the indispensable sources of pastoral rations, such as meat and dairy products, and used to be one of the means of transport by herders (Hund 2008). Furthermore, Mongolian horse was an essential and distinguished war-horse in history (Article: The Horse in Mongolian Culture 2018). For the Naadam of traditional festivals in Mongolia, horse racing is one of the entertaining activities for herds and regarded as the second most popular sporting events after wrestling (Davis 2010). So, the running speed of Mongolian horses has been one of the focuses of attention. Despite not the fastest horse in the world, people still endeavor to improve running speed of the Mongolian horse through unremitting consideration and breeding.

In order to discuss the genetic variation between Abaga horse and Wushen horse in Mongolian horse strains, we planned to analyze data of the entire genome with second-generation sequencing technology to seek out exercise-related single nucleotide polymorphism (SNP) locus of Mongolian horse and offer a reference for identification and improvement of Mongolian horse varieties.

## MATERIALS AND METHODS

### Experimental animals and sample preparation

We collected jugular blood of 4 Abaga horses in the Inner Mongolia Abaga County and 10 Wushen horses in the Inner Mongolia Ordos City Wushen County. Adhering to the manufacturer’s instructions for the extraction of DNA from the whole blood, genome was extracted by the AxyPrep blood genomic DNA kit. Then we used the NanoDrop ™ 1000 spectrophotometer and polyacrylamide gel electrophoresis to detect the concentration and integrity of the genome. The concentration of extracted DNA was between 26.4 ng/μL and 34.4 ng/μL for subsequent library construction.

### Library preparation and whole-genome sequencing

A TruSeq DNA Sample Prep Kit was used to construct a sequencing library. Whole-genome sequencing of the horses was performed using the Illumina HiSeq X™ Ten Sequencing System.

### Data quality control and comparison to reference genome

The raw data of 639,723,611,100 bp was sequenced from 14 samples. The inferior quality reads which have sequencing adapter, higher than 10% of N (base of uncertainty) content or inferior mass base (Q≤5) content of higher than 50% were filtered out by in-house Perl/Python scripts to achieve clean data of 630,535,376,400 bp. The Q20, Q30, error rate, GC content and other information of these data were counted by in-house Perl/Python scripts (Table 1). The sequencing reads were mapped to reference genomes (Ensembl release 82) by BWAmem (bwa-0.7.8) (Li 2013), which PCR and optical repetition of results were removed by using Picard (Broad Institute 2018). Statistics of mapping rate, average depth and coverage of the data after comparison were computed by in-house Perl/Python scripts (Table 2).

**Table 1.** Data volume and quality

| Sample | Sample Type | RawData | CleanData | EffectiveRat | ErrorRate | Q20 | Q30 | GC Content |
| --- | --- | --- | --- | --- | --- | --- | --- | --- |
| AB01 | Abaga Horse | 48,216,007,500 | 47,152,467,900 | 97.79 | 0.03 | 96.21 | 92.24 | 42.89 |
| AB02 | Abaga Horse | 43,047,445,800 | 42,340,444,800 | 98.36 | 0.03 | 96.17 | 92.1 | 42.65 |
| AB03 | Abaga Horse | 47,273,694,300 | 46,714,855,800 | 98.82 | 0.03 | 95.76 | 91.36 | 42.55 |
| AB04 | Abaga Horse | 48,416,284,200 | 47,906,212,800 | 98.95 | 0.03 | 95.96 | 91.47 | 42.13 |
| WS01 | Wushen Horse | 49,415,626,200 | 48,784,894,800 | 98.72 | 0.03 | 95.69 | 91.23 | 41.97 |
| WS02 | Wushen Horse | 53,074,008,900 | 52,417,962,000 | 98.76 | 0.03 | 96.09 | 92.19 | 42.35 |
| WS03 | Wushen Horse | 45,358,642,200 | 44,789,835,900 | 98.75 | 0.03 | 95.87 | 91.47 | 41.97 |
| WS04 | Wushen Horse | 47,681,665,800 | 47,139,254,400 | 98.86 | 0.03 | 95.94 | 91.6 | 41.66 |
| WS05 | Wushen Horse | 50,647,742,700 | 50,012,300,700 | 98.75 | 0.03 | 95.86 | 91.47 | 41.84 |
| WS06 | Wushen Horse | 45,072,226,800 | 44,633,740,200 | 99.03 | 0.03 | 95.79 | 91.61 | 42.19 |
| WS07 | Wushen Horse | 34,681,049,100 | 34,040,596,500 | 98.15 | 0.05 | 92.2 | 84.65 | 43.07 |
| WS08 | Wushen Horse | 45,212,322,600 | 44,350,535,700 | 98.09 | 0.04 | 92.41 | 84.97 | 43.02 |
| WS09 | Wushen Horse | 44,382,563,100 | 43,593,247,800 | 98.22 | 0.04 | 92.47 | 85.07 | 43.23 |
| WS10 | Wushen Horse | 37,244,331,900 | 36,659,027,100 | 98.43 | 0.04 | 92.91 | 85.77 | 43.12 |
| Average |  | 45,694,543,650 | 45,038,241,171 |  | 0.03 | 94.95 | 89.80 | 42.47 |
| Total |  | 639,723,611,100 | 630,535,376,400 | 98.56% |  |  |  |  |

**Table 2.** Data comparison

| Sample | Total Reads | Mapping<br>Rate(%) | Average<br>Depth | Coverage at<br>least 1X(%) | Coverage at<br>least 4X(%) | Coverage at<br>least 10X(%) |
| --- | --- | --- | --- | --- | --- | --- |
| AB01 | 315216417 | 98.55 | 17.46 | 99.59 | 99.32 | 93.57 |
| AB02 | 282990429 | 98.49 | 15.82 | 99.56 | 99.24 | 89.18 |
| AB03 | 312280472 | 98.49 | 16.93 | 99.52 | 98.98 | 90.17 |
| AB04 | 320340217 | 98.22 | 18.25 | 99.57 | 99.15 | 93.1 |
| WS01 | 326298749 | 98.6 | 17.63 | 99.54 | 99.25 | 94.36 |
| WS02 | 350423462 | 98.41 | 19.65 | 99.57 | 99.24 | 94.72 |
| WS03 | 299507485 | 98.63 | 16.45 | 99.51 | 99.16 | 92.2 |
| WS04 | 315038236 | 98.52 | 17.28 | 99.51 | 98.98 | 91.77 |
| WS05 | 334519623 | 98.65 | 18.19 | 99.54 | 99.24 | 95.43 |
| WS06 | 298331035 | 98.52 | 16.88 | 99.53 | 99.23 | 92.89 |
| WS07 | 227540381 | 97.88 | 12.85 | 99.56 | 98.73 | 68.52 |
| WS08 | 296433759 | 97.97 | 16.75 | 99.61 | 99.32 | 90.53 |
| WS09 | 291390679 | 98 | 16.44 | 99.51 | 99.18 | 88.97 |
| WS10 | 245088985 | 98.13 | 13.85 | 99.6 | 99.05 | 76.55 |
| Average | 301099994.9 | 98.36142857 | 16.745 | 99.55142857 | 99.14785714 | 89.42571429 |

### Single nucleotide polymorphism calling and annotation

SNP calling was performed using the GATK HaplotypeCaller (v3.5) (McKenna *et al.* 2010). In order to evaluate the reliability of the detected SNP sites and filter inferior quality SNP, we used SAMTools for SNP detection (Li 2011). Simultaneously, the dbSNP database 5,019,393 SNPs and 670K chip site information were downloaded. The data was used as training set, and the detected SNPs were evaluated and filtered by using the GATK VQSR process. The standard for the retention of the final site is the tranche value of 99 (Ti / Tv=2.02). Finally, the SNPs of equine population were filtered: GQ> 10, MAF> 0.05, call rate> 0.9 (Figure 1). The variants after filtering were annotated by ANNOVAR (v2016-02-01) (Wang *et al.* 2010) (Table 3).

**Figure 1.**
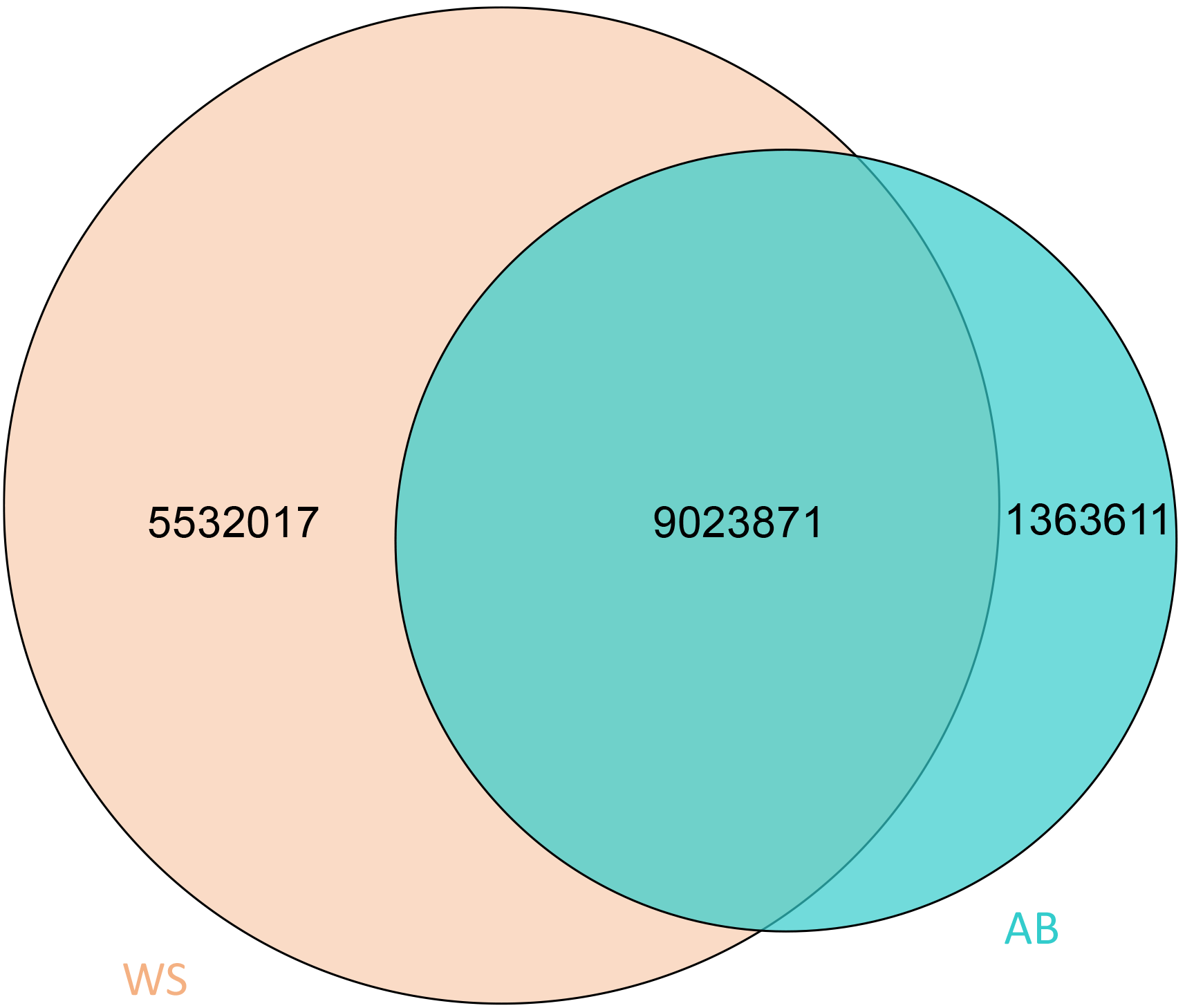
SNP following SNP calling. High quality SNPs were evaluated and indentified. AB, WS indicated Abaga horse and Wushen horse, respectively.

**Table 3.**
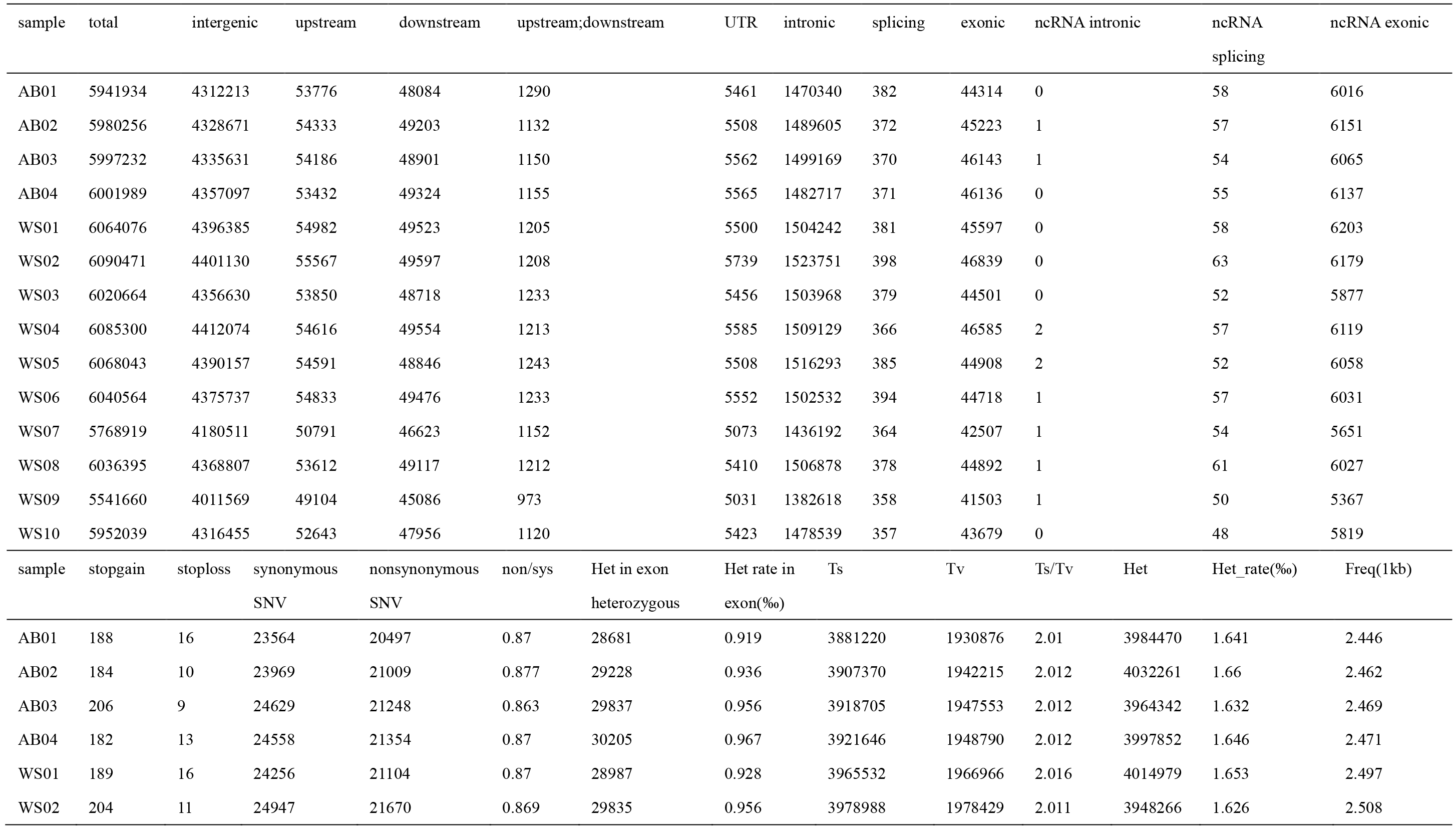
Single nucleotide polymorphism annotation

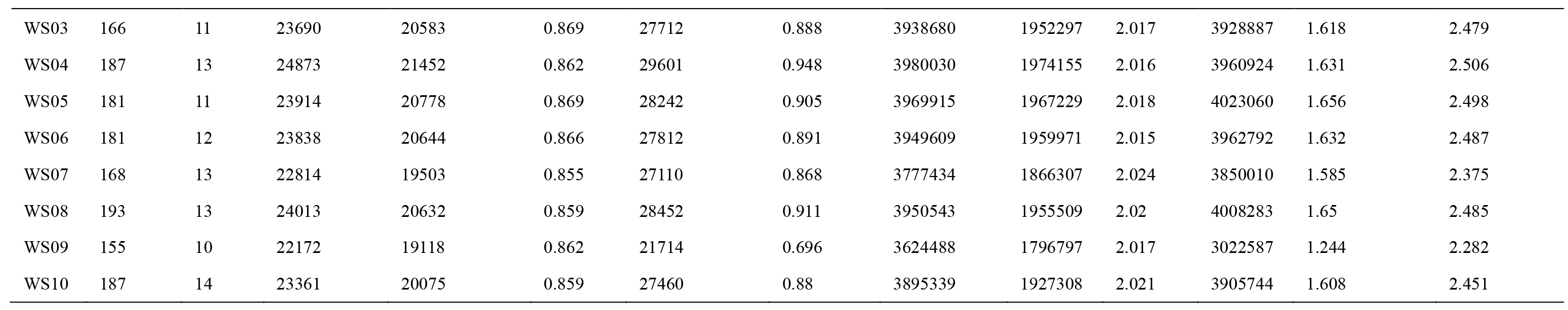

### Selective sweep analysis

To identify potential selective sweeps between Abaga horse (fast) and Wushen horse (slow), Pi log2(slow/fast) and FST was calculated together using VCFtools with a 20kb sliding window and a step size of 10 kb. Windows that contained less than 10 SNPs were excluded from further analysis. The windows that were simultaneously 1) in the top 5% of Z-transformed FST values and 2) in the bottom 5% Pi log2(slow/fast) were considered to be candidate selective regions in Abaga horse (Figure 2A; Table S1 https://doi.org/10.6084/m9.figshare.6289523.v1). The same applies to Wushen horse (Figure 2B; Table S2 https://doi.org/10.6084/m9.figshare.6289532.v1).

**Figure 2.**
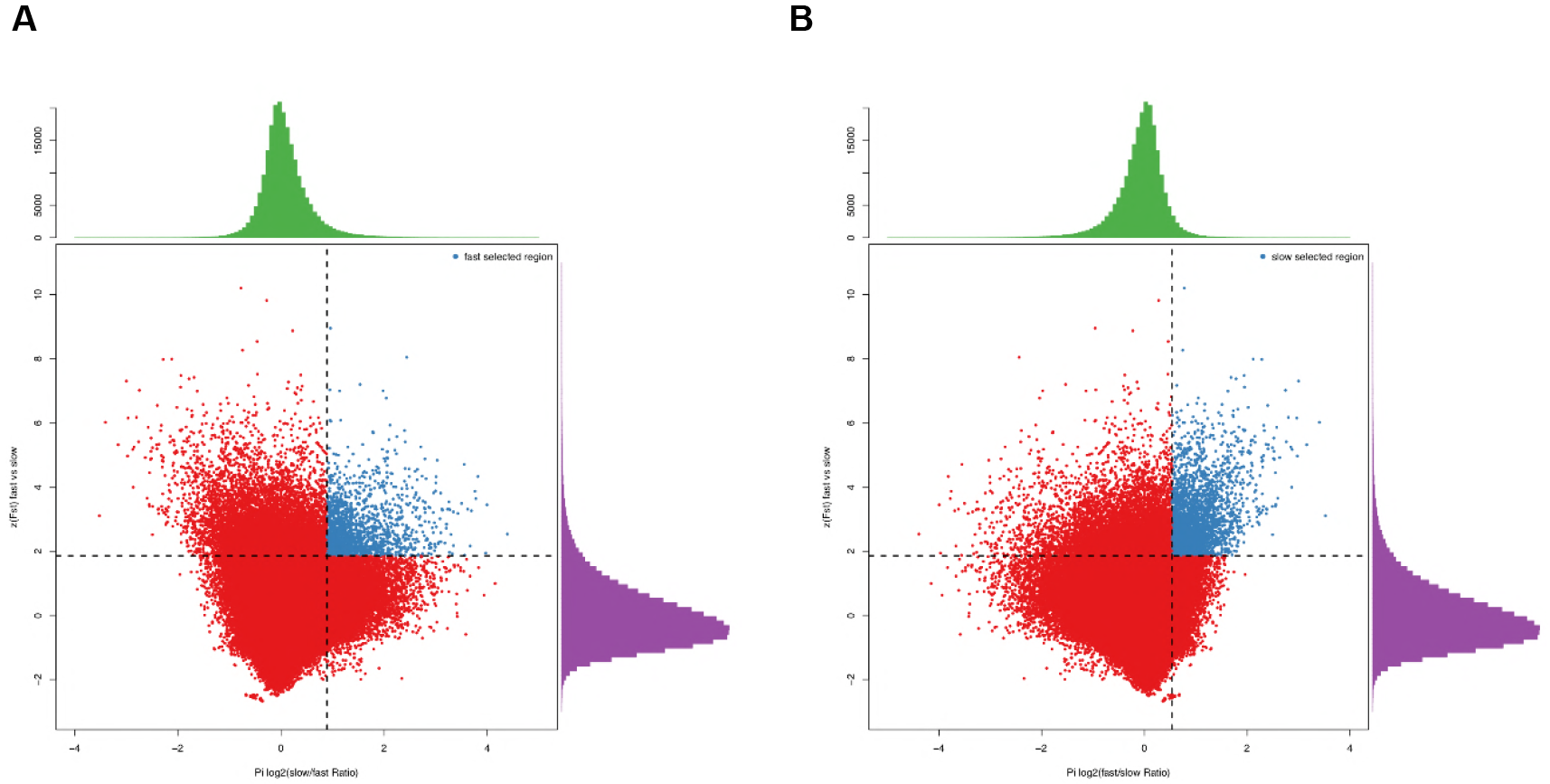
Identification of selected regions in Abaga horse and Wushen horse. To identify potential selective sweeps between Abaga horse (fast) and Wushen horse (slow), log2(πslow/πfast) and FST was calculated together using VCFtools with a 20kb sliding window and a step size of 10 kb. Windows that contained less than 10 SNPs were excluded from further analysis. The windows that were simultaneously 1) in the top 5% of Z-transformed FST values and 2) in the bottom 5% log2(πfast/πslow) were considered to be candidate selective regions in (A) Abaga horse and (B) Wushen horse.

### Statistic and advanced analysis of positively selected and candidate genes

We annotated the positively selected genes via GO (GOseq) to further screen out the major enriched functions (Young *et al.* 2010). The pathways which included these selected genes were enriched by KEGG (KOBAS) (Xie *et al.* 2011). Many positively selected genes which are in Abaga horse were analyzed further without overlapped genes between Abaga horse and Wushen horse.

## RESULTS AND DISCUSSION

### Related-clean data stated

We performed the entire genome second-generation sequencing for the whole blood of four Abaga horses (Figure 3A) and ten Wushen horses (Figure 3B) with the Illumina HiSeqX ten sequencing platform. The clean data of 630,535,376,400 bp (effective rate of data: 98.56%, error rate of data: 0.03%, mean of Q20: 94.95% and mean of Q30: 89.80%) was sequenced by filtration and 41.95G as the mean of clean data was generated in each sample (Table 1). Then, the data was mapped to the reference genome (Ensembl release 82) via BWAmem (Li 2013). Without PCR and optical repetition, the successful mapping rate of data was 98.36%. For 14 samples, the average sequencing depth was 16.75× coverage and average cover degree was 99.55% on reference sequences (Table 2).

**Figure 3.**
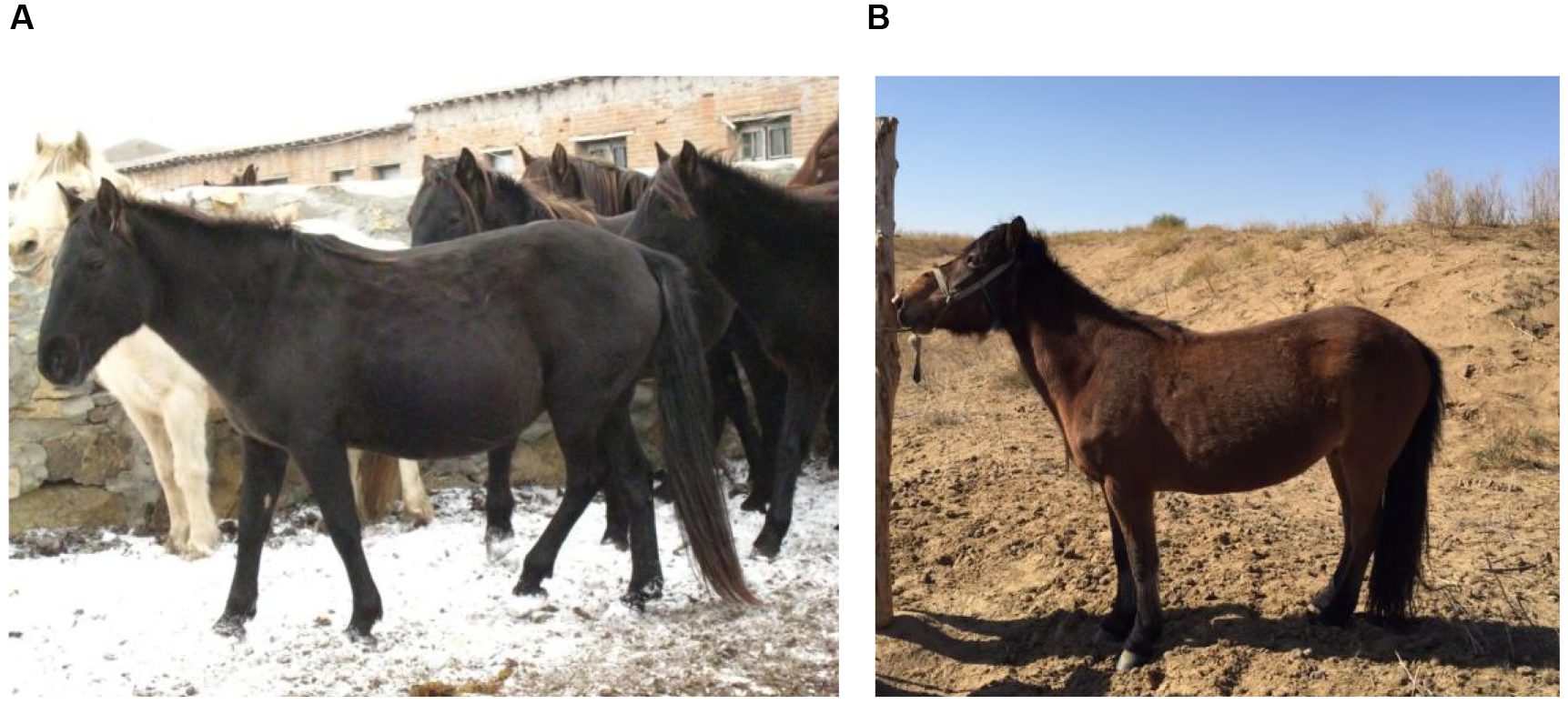
Actual Photos of Mongolian horses, (A) Abaga horse and (B) Wushen horse.

### Distribution of positive selection genes on chromosomes

Based on the data of SNP following SNP calling (Figure 1), we obtained the genes of significantly genetic differences by using F-statistics between Abaga horses and Wushen horses, and narrowed the above genes down to 479 and 943 positively selected genes combined with SNP polymorphism analysis in Abaga horses and Wushen horses, respectively (Figure 2A, B; Table S1, S2). We discovered that these selected genes were mainly distributed on chromosome 4, 7 and 10 in Abaga horses, and chromosome 1, 4, 8 and 16 in Wushen horses with the analysis of genes distribution on chromosome (Figure 4).

**Figure 4.**
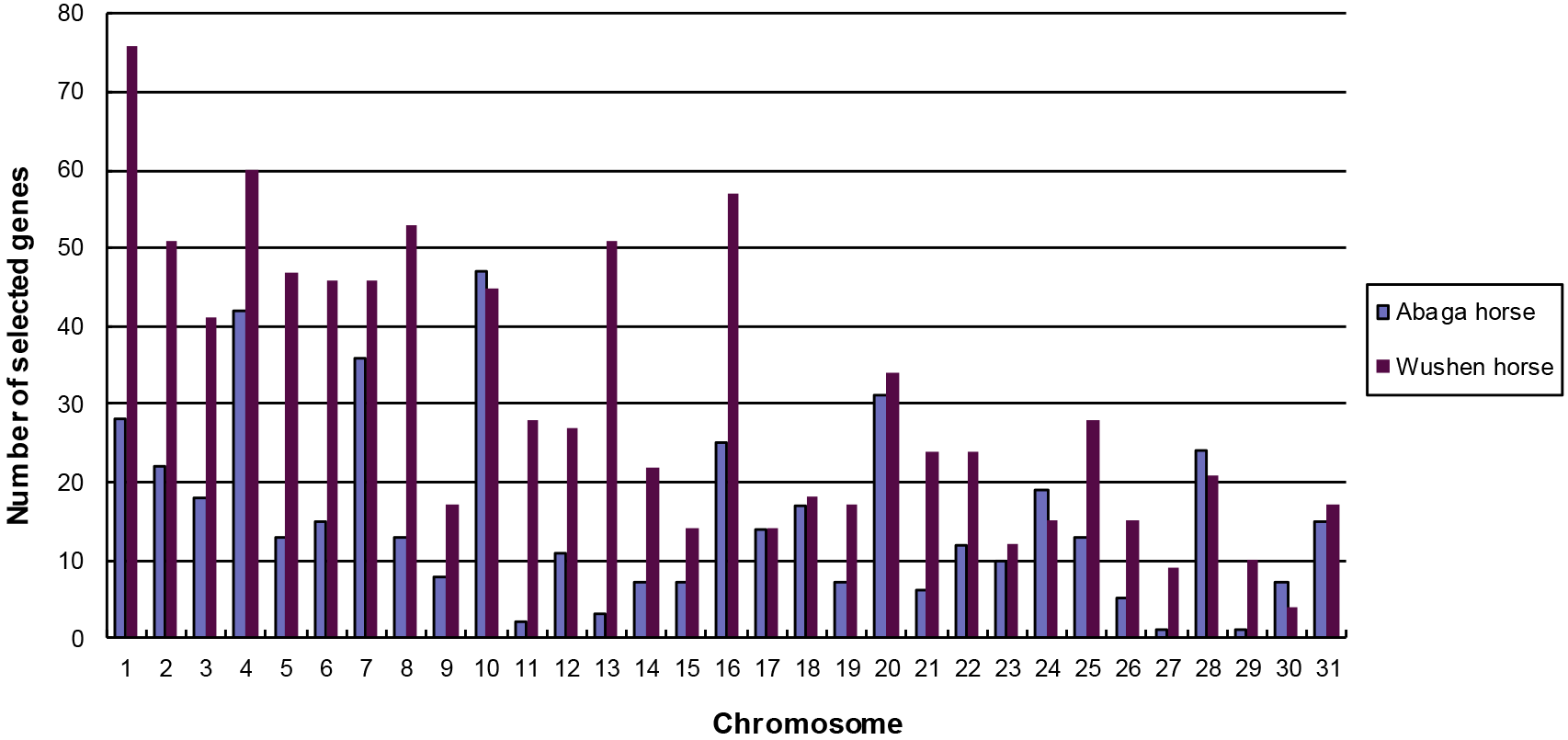
Distribution of selected genes on chromosomes.

Many genes of the positively selected 479 and 943 genes were enriched on chromosome 4, and the enrichment quantity of positively selected genes was secondary by the chromosome enrichment analysis both in Abaga horse and Wushen horse. In the statement of Schröder (Schröder *et al.* 2011), athletic performance-related genes were significantly enriched on chromosomes 4 and 12 of horses, which coincided with the different traits of running speed in our exploring direction. Possibly, we will take equine chromosome 4 as the exercise-related emphasis of scientific research.

### Gene Ontology, KEGG pathways and exercise-related genes

Above these positively selected genes were functional annotation by Gene Ontology (GO). The selected 479 genes of Abaga horse were mainly enriched in neuron part (GO:0097458), neuron projection (GO:0043005), regulation of membrane potential (GO:0045838), positive regulation of cell projection organization (GO:0031346), neuron-neuron synaptic transmission (GO:0007270), synaptic transmission, glutamatergic (GO:0035249), neurotransmitter secretion (GO:0007269), antigen processing and presentation (GO:0019882), telencephalon cell migration (GO:0022029) and forebrain cell migration (GO:0021885). The selected 943 genes of Wushen horse were mainly enriched in membrane part (GO:0044425), intrinsic component of membrane (GO:0031224), integral component of membrane (GO:0016021), cell projection (GO:0042995), neuron part (GO:0097458), neuron projection (GO:0043005), synapse (GO:0045202), cilium (GO:0005929) and cell projection assembly (GO:0030031) (Figure 5A, B).

**Figure 5.**
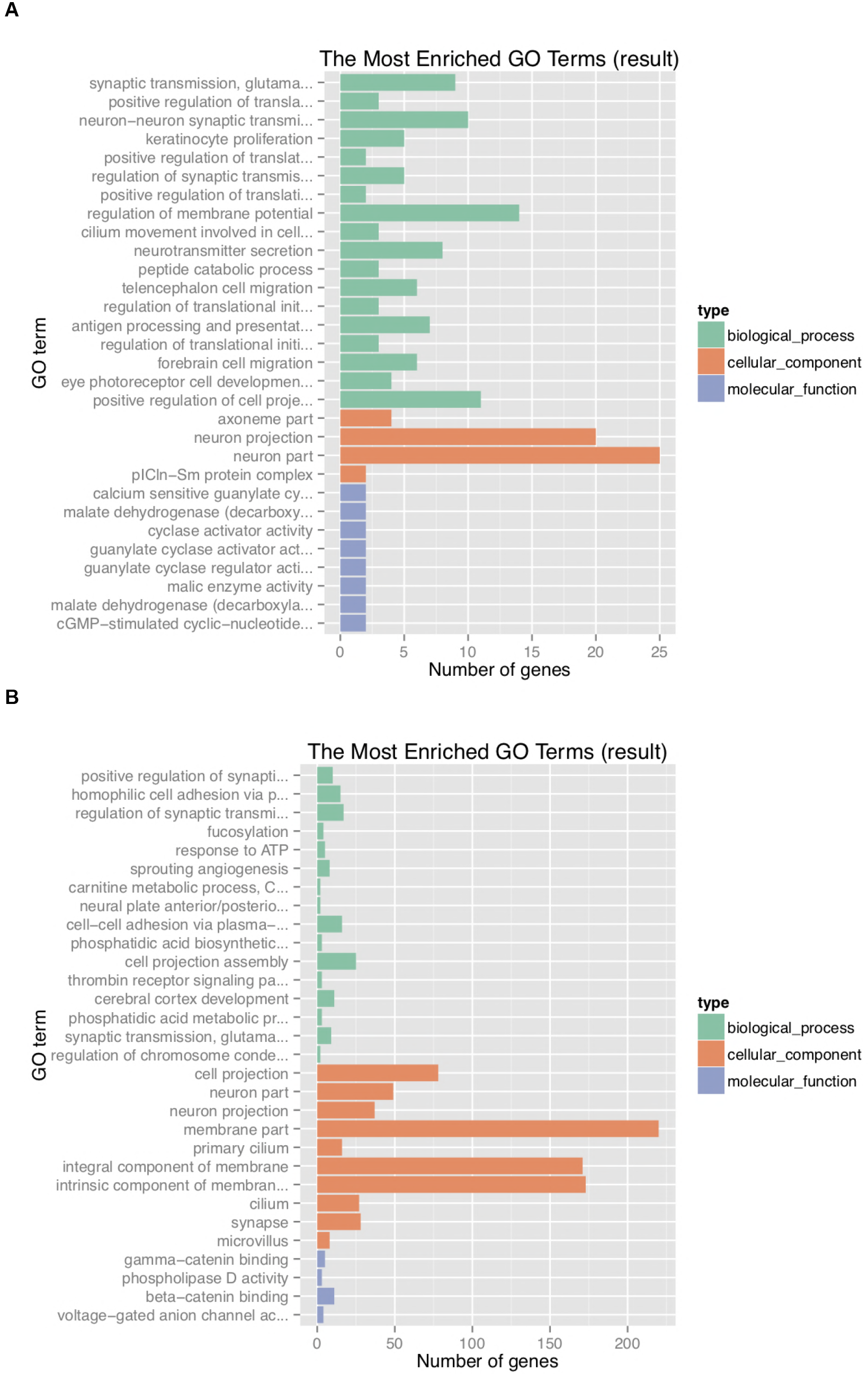
Function analysis based on Gene Ontology. The most enriched GO terms in (A) Abaga horse and (B) Wushen horse.

The athletic ability of the horse may be influenced not only by physiology, but also thought and motive. According to the previous studies, equine exercise-related genes included *DRD1-5*, *SLC6A4* and *BDNF*, the three genes functions were related to many neurological processes, involving motivation, pleasure, cognition, memory, learning, fine motor control, modulation of neuroendocrine signaling, adaptive ability of controlling emotions, supporting the survival of existing neurons, encouraging the growth and differentiation of new neurons and synapses (Momozawa *et al.* 2005; Bryan *et al.* 2007; Kulikova *et al.* 2007; Lippi *et al.* 2010). The GO analysis results of Abaga horse were also preferentially enriched in neuronal composition, neurotransmission and brain cell migration. These genes may allow Abaga horse to quickly observe and distinguish the surrounding during moving with high-speed, timely rectify status to respond to the various circumstances.

By pathways enrichment analysis of Kyoto Encyclopedia of Genes and Genomes (KEGG) with the positively selected 479 genes in Abaga horse and 943 genes in Wushen horse, the enriched pathways (P≤0.05) of Abaga horse included Propanoate metabolism, Viral myocarditis, Phototransduction, PI3K-Akt signaling pathway, Glycerolipid metabolism, Morphine addiction and mRNA surveillance pathway in which the pathway with the largest number of enriched genes (13 genes) was PI3K-Akt signaling pathway. As intracellular basal signaling pathways, PI3K-Akt signaling pathway involves lots of vital movement, such as exercise-induced physiologic hypertrophy (Shioi *et al.* 2002; Luo *et al.* 2005; Sagara *et al.* 2012; Song *et al.* 2015) and protecting mitochondria of skeletal muscle by aerobic endurance training (Liu *et al.* 2016), further explaining the excellent athletic performance of Abaga horse. Besides that, the enriched pathways (P≤0.05) of Wushen horse contained Base excision repair, Glutamatergic synapse, Endometrial cancer, Glycolysis / Gluconeogenesis, Propanoate metabolism and ABC transporters (Figure 6A, B).

**Figure 6.**
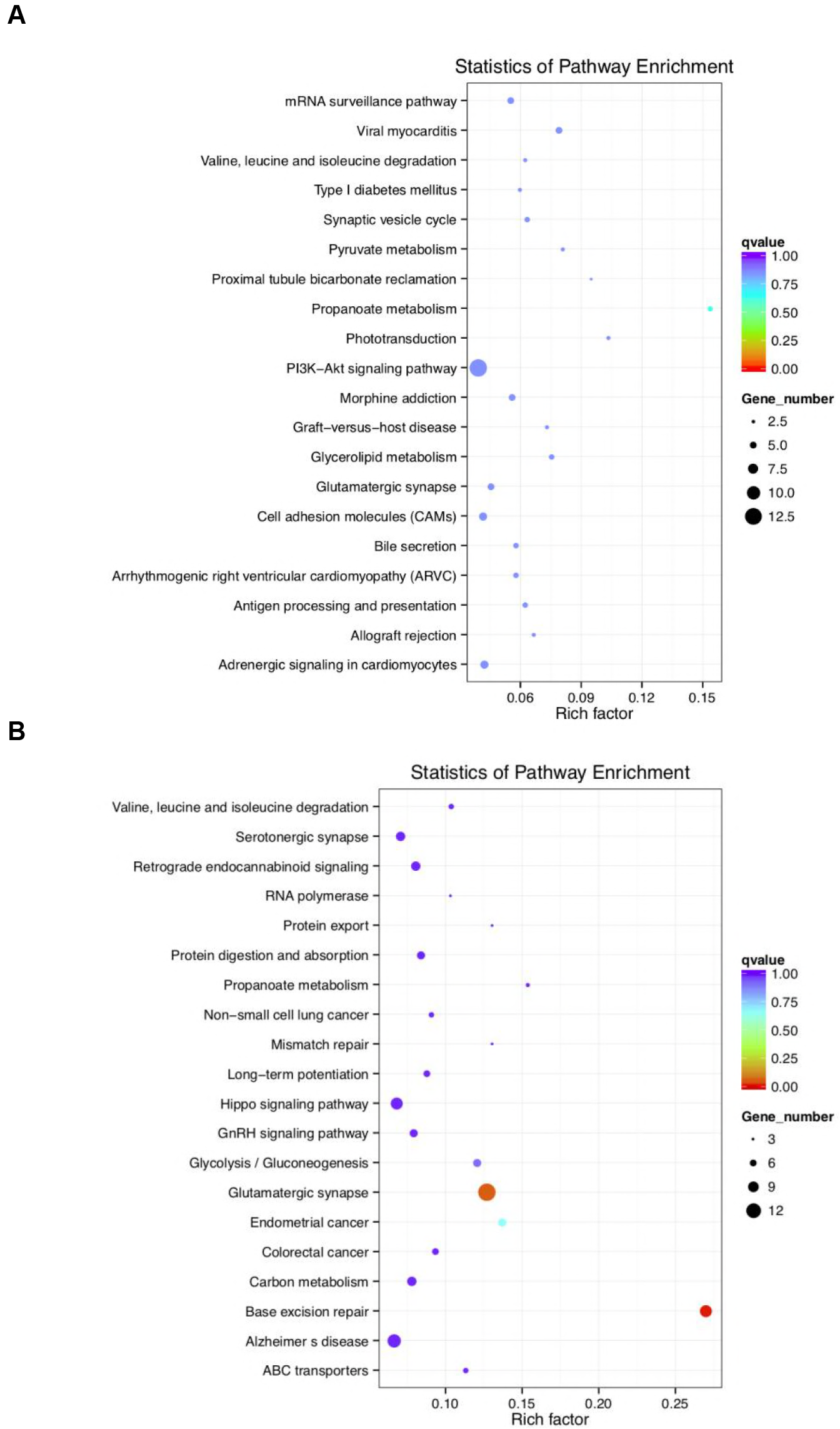
The KEGG pathway enrichment analysis. Top 20 of enriched pathways by statistics in (A) Abaga horse and (B) Wushen horse.

Further on SNPs, we analyzed functions of 440 genes of Abaga horse without 39 overlapped genes of positively selected genes between Abaga horse and Wushen horse. We focused on the enriched exercise-related pathways which referred to Metabolic pathways (Hill *et al.* 2010), Ras signaling pathway (Shioi *et al.* 2002; Xie *et al.* 2007), PI3K-Akt signaling pathway (Shioi *et al.* 2002; Luo *et al.* 2005; Sagara *et al.* 2012; Standard *et al.* 2014; Song *et al.* 2015; Liu *et al.* 2016; Vega *et al.* 2017), MAPK signaling pathway (Schröder *et al.* 2011), Hippo signaling pathway (Gabriel *et al.* 2016), Valine, leucine and isoleucine degradation (McGivney *et al.* 2010), Cardiac muscle contraction (Do *et al.* 2015), NF-kappa B signaling pathway (Kramer and Goodyear 2007), Arachidonic acid metabolism (Hill *et al.* 2010), Regulation of actin cytoskeleton (Hill *et al.* 2010; Schröder *et al.* 2011), Insulin signaling pathway (Gim *et al.* 2004; Hill *et al.* 2010) and Fatty acid metabolism (Gim *et al.* 2004; Hill *et al.* 2010) in the 440 positively selected genes of Abaga horse that distinguished from Wushen horse (Figure 7). These enriched pathways comprised some recurrent genes (Figure 7). Taking repeated genes as pivots, we speculated that the synergistic effect of pathways enabled faster running speed of Abaga horse compared with Wushen horse. But, in our studying, the enriched genes of positive selection were different from the previous studied genes in the above exercise-related pathways (Table 4), which indicated species-specific genes of positive selection in Abaga horse compared with other species (human, rat, mouse, leopard, thoroughbred horse etc.).

**Figure 7.**
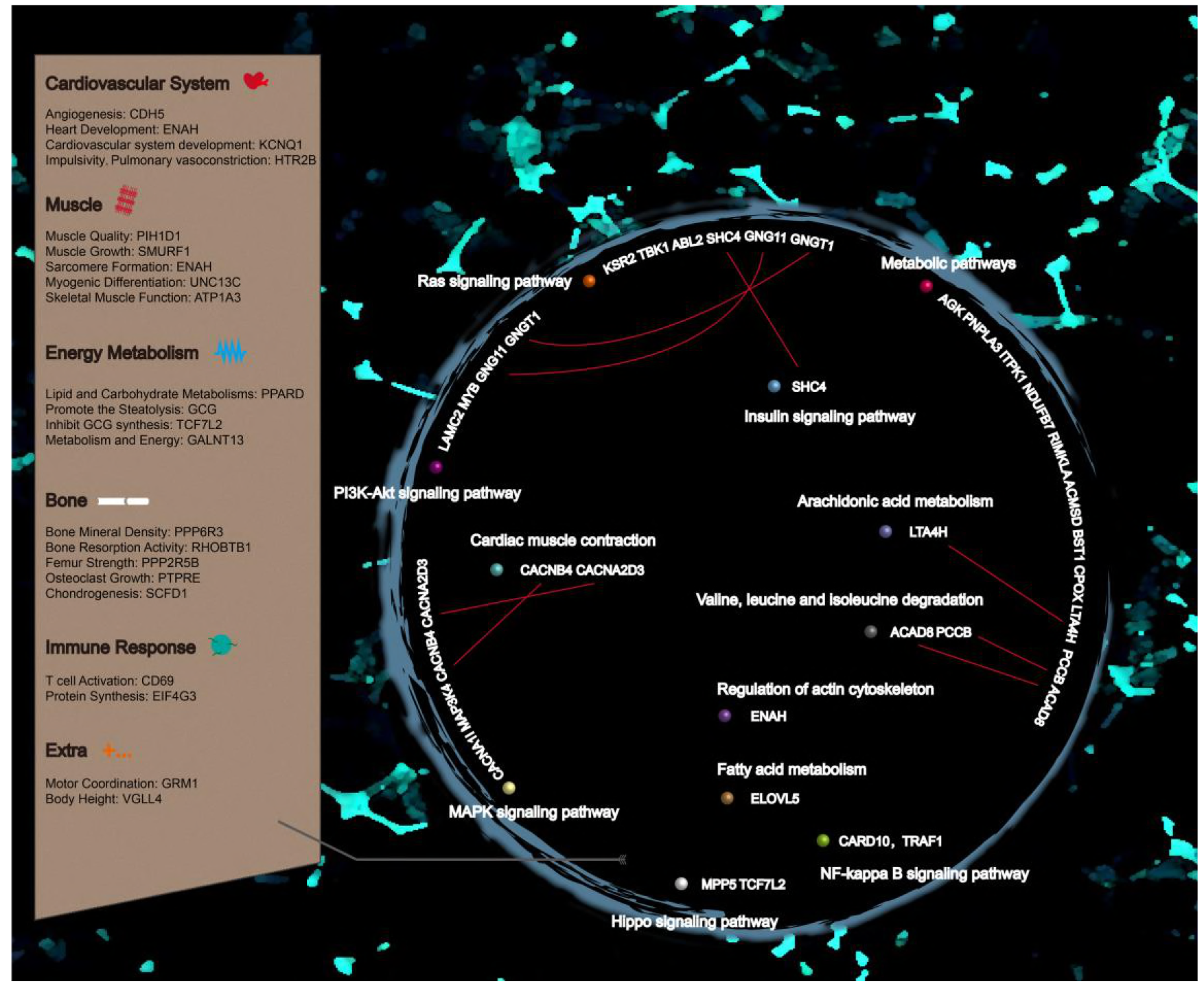
The exercise The exercise The exercise-related candidate genes and pathways of Abaga horse.

**Table 4.** Comparison of enriched genes in candidate pathways in Abaga horse and previous studies

| KEGG pathway | ID | Selected genes in Abaga horse | Selected genes or proteins in previous studies |
| --- | --- | --- | --- |
| Metabolic pathways | ecb01100 | *LTA4H AGK PNPLA3 ITPK1 NDUFB7 RIMKLA PCCB ACAD8 ACMSD BST1 CPOX* | *CYP51A1* |
| Ras signaling pathway | ecb04014 | *SHC4 GNG11 GNGT1 KSR2 TBK1 ABL2* | *APOA1 IGF-1 HRAS* |
| PI3K-Akt signaling pathway | ecb04151 | *LAMC2 GNG11 GNGT1 MYB* | PGC-1a IGF-1 IGF-1R<br>ErbB2 ErbB4 |
| MAPK signaling pathway | ecb04010 | *CACNA1I CACNA2D3 CACNB4 MAP3K4* | ERK AP-1 |
| Hippo signaling pathway | ecb04390 | *MPP5 TCF7L2* | *WWTR1 LATS2 TEAD YAP1 VGLL2 VGLL3 VGLL4* |
| Cardiac muscle contraction | ecb04260 | *CACNA2D3 CACNB4* | *CK-M* |
| NF-kappa B signaling pathway | ecb04064 | *CARD10 TRAF1* | MnSOD iNOS |
| Arachidonic acid metabolism | ecb00590 | *LTA4H* | *PTGS1* |
| Regulation of actin cytoskeleton | ecb04810 | *ENAH* | *GSN BDKRB2 CHRM MYLK ACTN3* |
| Insulin signaling pathway | ecb04910 | *SHC4* | *GYS1 PPARGC1A* |
| Fatty acid metabolism | ecb01212 | *ELOVL5* | *ADHFE1 SREBP2* |

According to the analysis of GO, KEGG and individual gene function, we subsequently put our interest in exercise-related genes of Abaga horse. Twenty-one genes may involve in exercise of Abaga horse while their functions embodied vasoconstriction (*HTR2B*) (Launay *et al.* 2002; Bevilacqua *et al.* 2010; Meira *et al.* 2014), angiogenesis (*CDH5*) (Sauteur *et al.* 2014), cardiac contraction (*KCNQ1*) (Jespersen *et al.* 2005; Brown *et al.* 2015; Pedersen *et al.* 2017), cardiac development and muscle structure (*ENAH*) (Franzini-Armstrong 1973; Benz *et al.* 2013), muscle growth (*PIH1D1, SMURF1*) (Inoue *et al.* 2010; Ponsuksili *et al.* 2014; Dalbo *et al.* 2013), myogenic differentiation (*UNC13C*) (Meyer *et al.* 2015; Langlois and Cowan 2017), skeletal muscle function (*ATP1A3*) (Aughey *et al.* 2007; Brashear *et al.* 2007), femur strength and bone mineral density (*PPP2R5B, PPP6R3*) (Alam *et al.* 2009; Medina-Gomez *et al.* 2017), osteoclast growth (*PTPRE, RHOBTB1*) (Chiusaroli *et al.* 2004; Song *et al.* 2014), chondrogenesis (SCFD) (DeLise *et al.* 2000; Hou *et al.* 2017), lipid and carbohydrate metabolism (*PPARD, GCG, TCF7L2, GALNT13*) (Yi *et al.* 2005; Bevilacqua *et al.* 2010; Park *et al.* 2012; Ahmetov and Fedotovskaya 2015; Giordano Attianese and Desvergne 2015; Ropka-Molik *et al.* 2017), exercise stress-induced response (*CD69, EIF4G3*) (Testi *et al.* 1989; Gradi *et al.* 1998; Cappelli *et al.* 2007; Morabito *et al.* 2016), exercise coordination (*GRM1*) (Conquet *et al.* 1994; Bossi *et al.* 2017) and height (*VGLL4*) (Gabriel *et al.* 2016). These genes of positive selection were presented simultaneously in Abaga horse, which may be a reason that it runs rapider than Wushen horse.

Counting on exercise-related genes of previous studies, the equine athletic performance is related to glucose metabolism, stress immune response, angiogenesis and muscle supply, insulin signal transduction, fat substrate application, muscle strength and the formation of bones and cartilage with growth (Gim *et al.* 2004; Hill *et al.* 2010; Park *et al.* 2012; Capomaccio *et al.* 2013; Kamm *et al.* 2013). We picked up exercise-related genes as candidate genes in positively selected genes, further, presented enriched KEGG pathways and functions with selected exercise-related genes (Figure 7). *HTR2B* (encoding 5-hydroxytryptamine receptor 2B), has been identified in the genome of Quarter horses of the racing line (Meira *et al.* 2014) and associated with impulsive behavior (Bevilacqua *et al.* 2010) and vasoconstriction (Launay *et al.* 2002). In zebrafish, vascular endothelial cadherin (encoded by *CDH5*) can promote elongation of the endothelial cell interface during angiogenesis (Sauteur *et al.* 2014). *KCNQ1* (encoding KvLQT1, a potassium channel protein) is related to exercise, and mutation of *KCNQ1* and *KCNE1* can casus susceptibility of sudden cardiac death (SCD) for horse (Jespersen *et al.* 2005; Brown *et al.* 2015; Pedersen *et al.* 2017). Mena (encoded by *ENAH*) which located in Z line that the borders of the sarcomere, VASP, and αII-Spectrin assemble cardiac multi-protein complexes to regulate cytoplasmic actin networks (Franzini-Armstrong 1973; Benz *et al.* 2013). *PIH1D1* (encoding the components of the apoptotic regulatory complex R2TP) is relevant to muscle mass (Inoue *et al.* 2010; Ponsuksili *et al.* 2014). Because E3 ubiquitin-protein ligase SMURF1 (encoded by *SMURF1*) function as negative regulators of myostatin pathway activity and myostatin is negative regulator of skeletal muscle mass, up-regulated expression of *SMURF1* may link to skeletal muscle growth following prolonged training (Dalbo *et al.* 2013). *UNC13C* is connected with differentiation of myoblast while integral myotubes originate in myoblast differentiation and raise the distinct muscle fiber types to build the complex skeletal muscle architecture for body movement, postural behavior and breathing (Meyer *et al.* 2015; Langlois and Cowan 2017). *ATP1A3* encodes subunit alpha-3 of sodium/potassium-transporting ATPase, which increased expression may be conducive to decrease fatigue after training (Brashear *et al.* 2007; Aughey *et al.* 2007). *PPARD* (encoding peroxisome proliferator-activated receptor delta) participate in regulation of energy metabolism, cell proliferation and differentiation, protection in stress conditions such as oxidative stress and inflammation and other important life activities (Giordano Attianese and Desvergne 2015). The antecedent studies have shown that Arabian horse will change the expression of *PPARD* and other genes of PPAR signaling pathway genes in skeletal muscle during exercise, and improve coefficient of utilization of fatty acids by energy conversion (Ropka-Molik *et al.* 2017). The up-regulated *PPARD* are also found after exercise in Thoroughbred horse (Park KD et al. 2012). So, we speculated that positively selected *PPARD* improved athletic ability by a similar mechanism in Abaga horse. Besides counter-regulatory hormone of insulin, *GCG* (encoding glucagon) is deemed to be involved in adipose metabolism and energy balance (Bevilacqua *et al.* 2010). Transcription factor 7-like 2 (encoded by *TCF7L2*) not only affects the metabolism of adipocytes by DNA methylation, but also activates the corresponding target genes through the Wnt signaling pathway to specifically inhibit glucagon synthesis in enteroendocrine cells (Yi *et al.* 2005). *GALNT13* may be involved in metabolic and energy pathways (Ahmetov and Fedotovskaya 2015).

Exercise has a great influence on the composition of the developing horse joints, the thickness of the hyaline cartilage of the adult horse, the calcified cartilage and subchondral bone (van de Lest *et al.* 2002; Tranquille *et al.* 2009). We found several genes associated with skeleton and cartilage development among candidate genes of Abaga horse. *PPP2R5B* and *PPP6R3* are closely related to femur strength in rats and bone mineral density in humans, respectively (Alam *et al.* 2009; Medina-Gomez *et al.* 2017). *PTPRE* encodes receptor-type tyrosine-protein phosphatase epsilon which is a positive regulator of osteoclast function (Chiusaroli *et al.* 2004). *RHOBTB1* is involved in osteoclast-mediated bone absorption activity (Song *et al.* 2014). Chondrogenesis demands transformation of chondrocytes from a simple mesenchymal condensation to cells with a highly enriched extracellular matrix (ECM) in the developing skeleton in which *SCFD1* plays an important role in the secretion of ECM protein during chondrogenesis (DeLise *et al.* 2000; Hou *et al.* 2017). So far there are no studies of association between these genes and the motor function of horses, but these skeleton- and cartilage-related genes provide new inspiration into the correlational research between ossature and exercise.

After exercise, the equine stress reaction will involve inflammation, cell signaling, and immune interactions (Capomaccio *et al.* 2013). Cell activation is the first step in the proliferation of immune cells, and CD69 is firstly detected in cell surface glycoproteins after activation (Testi *et al.* 1989). The low-to moderate-intensity aerobic trekking induces activation of CD69 T cell and promotes anti-stress effects on the oxidative balance and the high-altitude-induced injury of the immune responses among women (Morabito *et al.* 2016). *EIF4G3* encodes eukaryotic translation initiation factor 4 gamma 3 which is indispensable for triggering protein synthesis and is thought to be involved in exercise stress-induced response in horses (Gradi *et al.* 1998; Cappelli *et al.* 2007). We hypothesized that these genes may be involved in the ability of Abaga horses to enhance certain diseases resistance through exercise, but more data and experiments are needed to verify.

*GRM1* encodes metabotropic glutamate receptor 1, which deficiency can lead to serious deficits of motor coordination and spatial learning in mice (Conquet *et al.* 1994; Bossi *et al.* 2017). The effectors of Hippo signal pathway regulate several motor-related genes and adaptations while *VGLL4* is Hippo-signal-related to body height (Gabriel *et al.* 2016). These exercise-related genes were positively selected in Abaga horse, indicating that Abaga horse has exercise-related genetic potential compared with Wushen horse.

In conclusion, we analyzed the genomic data of Abaga horse and Wushen horse by sequencing. We uncovered that most of the positively selected genes, particularly exercise-related, of Abaga horse and Wushen horse were concentrated on chromosome 4, which implied that the chromosome 4 may be associated with the evolution of the Mongolian horse, and athletic performance may be the future research direction. The positively selected genes of Abaga horse were enriched in exercise-related pathways that were different from some selected genes of other horses or species, suggesting that the Abaga horse may have exclusively physiological mechanism for the motorial process. Twenty-one exercise-related genes were detected, which provided molecular genetic basis for further research on athletic performance and breeding of Mongolian horse.

## ACKNOWLEDGMENTS

We thank Y.R. Zhang, K.F. Wu, S.Y. Wang, Z.Z. Liu, W. Wei, P.W. Shi and C.H. Cao for sampling assistance.

## SUPPLEMENTAL MATERIALS

**Table S1** The genes of the selective region in Abaga horse.

**Table S2** The genes of the selective region in Wushen horse.

## LITERATURE CITED

Ahmetov I.I., Fedotovskaya O.N., 2015. Current Progress in Sports Genomics. Adv Clin Chem. 70:247–314.

Article: The Horse in Mongolian Culture, 2018. https://www.amnh.org/explore/science-bulletins/bio/documentaries/the-last-wild-horse-the-return-of-takhi-to-mongolia/article-the-horse-in-mongolian-culture. Accessed 10 Jan 2018.

Alam I., Sun Q., Koller D.L., Liu L., Liu Y., Edenberg H.J., Li J., Foroud T., Turner C. H., 2009. Differentially expressed genes strongly correlated with femur strength in rats. Genomics 94:257–62.

Aughey R.J., Murphy K.T., Clark S.A., Garnham A.P., Snow R.J., Cameron-Smith D., Hawley J.A., McKenna M.J., 2007. Muscle Na+-K+-ATPase activity and isoform adaptations to intense interval exercise and training in well-trained athletes. J Appl Physiol (1985). 103:39–47.

Benz P.M, Merkel C.J, Offner K., AbeBer M., Ullrich M., Fischer T., Bayer B., Wagner H., Gambaryan S., Ursitti J.A., Adham I.M., Linke W.A., Feller S.M., Fleming I., Renné T., Frantz S., Unger A., Schuh K., 2013. Mena/VASP and aII-Spectrin complexes regulate cytoplasmic actin networks in cardiomyocytes and protect from conduction abnormalities and dilated cardiomyopathy. Cell Commun Signal. 11:56.

Bevilacqua L., Doly S., Kaprio J., Yuan Q., Tikkanen R., Paunio T., Zhou Z., Wedenoja J., Maroteaux L., Diaz S., Belmer A., Hodgkinson C.A., Dell’osso L., Suvisaari J., Coccaro E., Rose R.J., Peltonen L., Virkkunen M., Goldman D., 2010. A population-specific HTR2B stop codon predisposes to severe impulsivity. Nature. 468:1061–6.

Bossi S., Musante I., Bonfiglio T., Bonifacino T., Emionite L., Cerminara M., Cervetto C., Marcoli M., Bonanno G., Ravazzolo R., Pittaluga A., Puliti A., 2017. Genetic inactivation of mGlu5 receptor improves motor coordination in the Grm1^crv4^ mouse model of SCAR13 ataxia. Neurobiol Dis. 109:44–53.

Brashear A., Dobyns W.B., de Carvalho Aguiar P., Borg M., Frijns C.J., Gollamudi S., Green A., Guimaraes J., Haake B.C., Klein C., Linazasoro G., Münchau A., Raymond D., Riley D., Saunders-Pullman R., Tijssen M.A., Webb D., Zaremba J., Bressman S.B., Ozelius L.J., 2007. The phenotypic spectrum of rapid-onset dystonia-parkinsonism (RDP) and mutations in the ATP1A3 gene. Brain. 130:828–35.

Broad Institute. Picard Tools - By Broad Institute. http://broadinstitute.github.io/picard. Accessed 11 Jan 2018.

Brown W.M., 2015. Exercise-associated DNA methylation change in skeletal muscle and the importance of imprinted genes: a bioinformatics meta-analysis. Br J Sports Med. 49:1567–78.

Bryan A., Hutchison K.E., Seals D.R., Allen D.L., 2007. A transdisciplinary model integrating genetic, physiological, and psychological correlates of voluntary exercise. Health Psychol. 26:30–9.

Capomaccio S., Vitulo N., Verini-Supplizi A., Barcaccia G., Albiero A., D’Angelo M., Campagna D., Valle G., Felicetti M., Silvestrelli M., Cappelli K., 2013. RNA sequencing of the exercise transcriptome in equine athletes. PLoS One. 8:e83504.

Cappelli K., Verini-Supplizi A., Capomaccio S., Silvestrelli M., 2007. Analysis of peripheral blood mononuclear cells gene expression in endurance horses by cDNA-AFLP technique. Res Vet Sci. 82:335–43.

China National Commission of Animal Genetic Resources., 2011. Animal Genetic Resources in China: Horse, Donkeys, Camels. China Agriculture Press, Beijing, pp. 28–37.

Chiusaroli R., Knobler H., Luxenburg C., Sanjay A., Granot-Attas S., Tiran Z., Miyazaki T., Harmelin A., Baron R., Elson A., 2004. Tyrosine phosphatase epsilon is a positive regulator of osteoclast function in vitro and in vivo. Mol Biol Cell. 15:234–44.

Conquet F., Bashir Z., Davies C.H., Daniel H., Ferraguti F., Bordi F., Franz-Bacon K., Reggiani A., Matarese V., Condé F., Collingridge G.L., Crépel F., 1994. Motor deficit and impairment of synaptic plasticity in mice lacking mGluR1. Nature. 372:237–43.

Dalbo V.J., Roberts M.D., Hassell S., Kerksick C.M. 2013. Effects of pre-exercise feeding on serum hormone concentrations and biomarkers of myostatin and ubiquitin proteasome pathway activity. Eur J Nutr. 52:477–87.

Davis M., 2010. When Things Get Dark: A Mongolian Winter’s Tale. St. Martin’s Press, NY, pp. 169.

DeLise A.M., Fischer L., Tuan R.S., 2000. C Cellular interactions and signaling in cartilage development. Osteoarthritis Cartilage. 8:309–34.

Do K.T., Cho H.W., Badrinath N., Park J.W., Choi J.Y., Chung Y.H., Lee H.K., Song K.D., Cho B.W., 2015. Molecular Characterization and Expression Analysis of Creatine Kinase Muscle (CK-M) Gene in Horse. Asian-Australas J Anim Sci. 28:1680–5.

Doan R., Cohen N.D., Sawyer J., Ghaffari N., Johnson C.D., Dindot S.V., 2012. Whole-genome sequencing and genetic variant analysis of a Quarter Horse mare. BMC Genomics. 13:78.

Dugarjaviin M., 2009. Horse in China. Hong Kong Cultural/China Horstry Publishing CO., Ltd, HK.

Elisabeth Y., 2011. The Mongolian Horse and Horseman. In: SIT Graduate Institute/SIT Study Abroad, SIT Digital Collections. http://digitalcollections.sit.edu/cgi/viewcontent.cgi?article=2074&context=ispcollection. Accessed 11 Jan 2018.

Franzini-Armstrong C., 1973. The structure of a simple Z line. J Cell Biol. 58:630–42.

Gabriel B.M., Hamilton D.L., Tremblay A.M., Wackerhage H., 2016. The Hippo signal transduction network for exercise physiologists. J Appl Physiol. (1985) 120:1105–17.

Gim J.A., Ayarpadikannan S., Eo J., Kwon Y.J., Choi Y., Lee H.K., Park K.D., Yang Y.M., Cho B.W., Kim H.S., 2014. Transcriptional expression changes of glucose metabolism genes after exercise in thoroughbred horses. Gene. 547: 152–8.

Giordano Attianese G.M., Desvergne B., 2015. Integrative and systemic approaches for evaluating PPARβ/δ (PPARD) function. Nucl Recept Signal. 13:e001.

Gradi A., Imataka H., Svitkin Y.V., Rom E., Raught B., Morino S., Sonenberg N., 1998. A novel functional human eukaryotic translation initiation factor 4G. Mol Cell Biol. 18:334–42.

Hill E.W., Gu J., McGivney B.A., MacHugh D.E., 2010. Targets of selection in the Thoroughbred genome contain exercise-relevant gene SNPs associated with elite racecourse performance. Anim Genet 41 Suppl. 2:56–63.

Hou N., Yang Y., Scott I.C., Lou X., 2017. The Sec domain protein Scfd1 facilitates trafficking of ECM components during chondrogenesis. Dev Biol. 421:8–15.

Hund A. 2008. The Stallion’s Mane The Next Generation of Horses in Mongolia. Parasite Immunology. 20:73–80.

Inoue M., Saeki M., Egusa H., Niwa H., Kamisaki Y., 2010. PIH1D1, a subunit of R2TP complex, inhibits doxorubicin-induced apoptosis. Biochem Biophys Res Commun. 403:340–4.

Jespersen T., Grunnet M., Olesen S.P., 2005. The KCNQ1 Potassium Channel: From Gene to Physiological Function. Physiology (Bethesda). 20:408–16.

Kamm J.L., Frisbie D.D., McIlwraith C.W., Orr K.E., 2013. Gene biomarkers in peripheral white blood cells of horses with experimentally induced osteoarthritis. Am J Vet Res. 74:115–21.

Kramer H.F., Goodyear L.J., 2007. Exercise, MAPK, and NF-kappaB signaling in skeletal muscle. J Appl Physiol (1985). 103:388–95.

Kulikova M.A., Maliuchenko N.V., Timofeeva M.A., Shleptsova V.A., Tschegol’kova IuA., Vediakov A.M., Tonevitskiĭ A.G., 2007. Polymorphisms of the main genes of neurotransmitter systems: I. the dopaminergic system. Fiziol Cheloveka. 33:105–12.

Langlois S., Cowan K.N., 2017. Regulation of Skeletal Muscle Myoblast Differentiation and Proliferation by Pannexins. Adv Exp Med Biol. 925:57–73.

Launay J.M., Hervé P., Peoc’h K., Tournois C., Callebert J., Nebigil C.G., Etienne N., Drouet L., Humbert M., Simonneau G., Maroteaux L., 2002. Function of the serotonin 5-hydroxytryptamine 2B receptor in pulmonary hypertension. Nat Med. 8:1129–35.

Li H., 2011. A statistical framework for SNP calling, mutation discovery, association mapping and population genetical parameter estimation from sequencing data. Bioinformatics. 27:2987–93.

Li H., 2013. Aligning sequence reads, clone sequences and assembly contigs with BWA-MEM. https://arxiv.org/abs/1303.3997. Accessed 26 Dec 2016.

Li L.F., Guan W.J., Hua Y., Bai X.J., Ma Y.H., 2009. Establishment and characterization of a fibroblast cell line from the Mongolian horse. In Vitro Cell Dev Biol Anim. 45:311–6.

Lippi G., Longo UG., Maffulli N., 2010, Genetics and sports. Br Med Bull, 93:27–47,

Liu S.D., Zhang Y.Q., Cao J., 2016. The influence of the aerobic endurance training on the skeletal muscular mitochondria function and PI3K-Akt protein expression. Zhongguo Ying Yong Sheng Li Xue Za Zhi. 32:55–8.

Luo J., McMullen J.R., Sobkiw C.L., Zhang L., Dorfman A.L., Sherwood M.C., Logsdon M.N., Horner J.W., DePinho R.A., Izumo S., Cantley L.C., 2005. Class IA Phosphoinositide 3-Kinase Regulates Heart Size and Physiological Cardiac Hypertrophy. Mol Cell Biol. 25:9491–502.

McGivney B.A., McGettigan P.A., Browne J.A., Evans A.C., Fonseca R.G., Loftus B.J., Lohan A., MacHugh D.E., Murphy B.A., Katz L.M., Hill E.W., 2010. Characterization of the equine skeletal muscle transcriptome identifies novel functional responses to exercise training. BMC Genomics. 11:398.

McKenna A., Hanna M., Banks E., Sivachenko A., Cibulskis K., Kernytsky A., Garimella K., Altshuler D., Gabriel S., Daly M., DePristo M.A., 2010. The Genome Analysis Toolkit: a MapReduce framework for analyzing next-generation DNA sequencing data. Genome Res. 20:1297–303.

Medina-Gomez C., Kemp J.P., Dimou N.L., Kreiner E., Chesi A., Zemel B.S., Bønnelykke K., Boer C.G., Ahluwalia T.S., Bisgaard H., Evangelou E., Heppe D.H.M., Bonewald L.F., Gorski J.P., Ghanbari M., Demissie S., Duque G., Maurano M.T., Kiel D.P., Hsu Y.H., C. J. van der Eerden B., Ackert-Bicknell C., Reppe S., Gautvik K.M., Raastad T., Karasik D., van de Peppel J., Jaddoe V.W.V., Uitterlinden A.G., Tobias J.H., Grant S.F.A., Bagos P.G., Evans D.M., Rivadeneira F., 2017. Bivariate genome-wide association meta-analysis of pediatric musculoskeletal traits reveals pleiotropic effects at the SREBF1/TOM1L2 locus. Nat Commun. 8:121.

Meira C.T., Curi R.A., Farah M.M., de Oliveira H.N., Béltran N.A., Silva J.A. 2nd., 2014. Prospection of genomic regions divergently selected in racing line of Quarter Horses in relation to cutting line. Animal. 8:1754–64.

Meyer S.U., Krebs S., Thirion C., Blum H., Krause S., Pfaffl M.W., 2015. Tumor Necrosis Factor Alpha and Insulin-Like Growth Factor 1 Induced Modifications of the Gene Expression Kinetics of Differentiating Skeletal Muscle Cells. PLoS One. 10:e0139520.

Momozawa Y., Takeuchi Y., Kusunose R., Kikusui T., Mori Y., 2005. Association between equine temperament and polymorphisms in dopamine D4 receptor gene. Mamm Genome. 16:538–44.

Morabito C., Lanuti P., Caprara G.A., Guarnieri S., Verratti V, Ricci G., Catizone A., Marchisio M., Fanò-Illic G., Mariggiò M.A., 2016. Responses of peripheral blood mononuclear cells to moderate exercise and hypoxia. Scand J Med Sci Sports. 26:1188–99.

Park K.D., Park J., Ko J., Kim B.C., Kim H.S., Ahn K., Do K.T., Choi H., Kim H.M., Song S., Lee S., Jho S., Kong H.S., Yang Y.M., Jhun B.H., Kim C., Kim T.H., Hwang S., Bhak J., Lee H.K., Cho B.W., 2012. Whole transcriptome analyses of six thoroughbred horses before and after exercise using RNA-Seq. BMC Genomics. 13:473.

Pedersen P.J., Thomsen K.B., Flak J.B., Tejada M.A., Hauser F., Trachsel D., Buhl R., Kalbfleisch T., DePriest M.S., MacLeod J.N., Calloe K., Klaerke D.A., 2017. Molecular cloning and functional expression of the K^+^, channel KV7.1 and the regulatory subunit KCNE1 from equine myocardium. Res Vet Sci. 113:79–86.

Ponsuksili S., Murani E., Trakooljul N., Schwerin M., Wimmers K., 2014. Discovery of candidate genes for muscle traits based on GWAS supported by eQTL-analysis. Int J Biol Sci. 10:327–37.

Ropka-Molik K., Stefaniuk-Szmukier M., Z Ukowski K., Piókowska K., Bugno-Poniewierska M., 2017. Exercise-induced modification of the skeletal muscle transcriptome in Arabian horses. Physiol Genomics. 49:318–326.

Sagara S., Osanai T., Itoh T., Izumiyama K., Shibutani S., Hanada K., Yokoyama H., Yamamoto Y., Yokota T., Tomita H., Magota K., Okumura K., 2012. Overexpression of coupling factor 6 attenuates exercise-induced physiological cardiac hypertrophy by inhibiting PI3K/Akt signaling in mice. J Hypertens. 30:778–86.

Sauteur L., Krudewig A., Herwig L., Ehrenfeuchter N., Lenard A., Affolter M., Belting H.G., 2014. Cdh5/VE-cadherin promotes endothelial cell interface elongation via cortical actin polymerization during angiogenic sprouting. Cell Rep. 9:504–13.

Schröder W., Klostermann A., Distl O., 2011. Candidate genes for physical performance in the horse. Vet J. 190:39–48.

Shioi T., McMullen J.R., Kang P.M., Douglas P.S., Obata T., Franke T.F., Cantley L.C., Izumo S., 2002. Akt/protein kinase B promotes organ growth in transgenic mice. Mol Cell Biol. 22:2799–809.

Song H.K., Kim J., Lee J.S., Nho K.J., Jeong H.C., Kim J., Ahn Y., Park W.J., Kim D. H., 2015. Pik3ip1 modulates cardiac hypertrophy by inhibiting PI3K pathway. PLoS One. 10:e0122251.

Song R., Gu J., Liu X., Zhu J., Wang Q., Gao Q., Zhang J., Cheng L., Tong X., Qi X., Yuan Y., Liu Z., 2014. Inhibition of osteoclast bone resorption activity through osteoprotegerin-induced damage of the sealing zone. Int J Mol Med. 34:856–62.

Standard J., Jiang Y., Yu M., Su X., Zhao Z., Xu J., Chen J., King B., Lu L., Tomich J., Baybutt R., Wang W., 2014. Reduced signaling of PI3K-Akt and RAS-MAPK pathways are the key targets for weight loss-induced cancer prevention by dietary calorie restriction and/or physical activity. J Nutr Biochem. 25: 1317–1323.

Testi R., Phillips J.H., Lanier L.L., 1989. Leu 23 induction as an early marker of functional CD3/T cell antigen receptor triggering. Requirement for receptor cross-linking, prolonged elevation of intracellular [Ca++] and stimulation of protein kinase C. J Immunol. 142:1854–60.

Tranquille C.A., Blunden A.S., Dyson S.J., Parkin T.D., Goodship A.E., Murray R.C., 2009. Effect of exercise on thicknesses of mature hyaline cartilage, calcified cartilage, and subchondral bone of equine tarsi. Am J Vet Res. 70:1477–83.

van de Lest C.H., Brama P.A., Van Weeren P.R., 2002. The influence of exercise on the composition of developing equine joints. Biorheology. 39:183–91.

Vega R.B., Konhilas J.P., Kelly D.P., Leinwand L.A., 2017. Molecular Mechanisms Underlying Cardiac Adaptation to Exercise. Cell Metab. 25:1012–1026.

Wang K., Li M., Hakonarson H., 2010. ANNOVAR: functional annotation of genetic variants from high-throughput sequencing data. Nucleic Acids Res. 38:e164.

Xie C., Mao X., Huang J., Ding Y, Wu J., Dong S., Kong L., Gao G., Li C.Y., Wei L., 2011. KOBAS 2.0: a web server for annotation and identification of enriched pathways and diseases. Nucleic Acids Res. 39:W316–22.

Xie L., Jiang Y., Ouyang P., Chen J., Doan H., Herndon B., Sylvester J.E., Zhang K., Molteni A., Reichle M., Zhang R., Haub M.D., Baybutt R.C., Wang W., 2007. Effects of dietary calorie restriction or exercise on the PI3K and Ras signaling pathways in the skin of mice. J Biol Chem. 282:28025–35.

Yi F., Brubaker P.L., Jin T., 2005. TCF-4 Mediates Cell Type-specific Regulation of Proglucagon Gene Expression by β-Catenin and Glycogen Synthase Kinase-3β;. J Biol Chem. 280:1457–64.

Young M.D., Wakefield M.J., Smyth G.K., Oshlack A., 2010. Gene ontology analysis for RNA-seq: accounting for selection bias. Genome Biol. 11:R14.

